# Transcriptional program of memory B cell activation, broadly binding anti-influenza antibodies, and bystander activation after vaccination revealed by single-cell transcriptomics

**DOI:** 10.1101/709337

**Authors:** Felix Horns, Cornelia L. Dekker, Stephen R. Quake

**Author notes:** Correspondence to (S.R.Q.).

## Abstract

Antibody memory protects humans from many diseases. Protective antibody memory responses require activation of transcriptional programs, cell proliferation, and production of antigen-specific antibodies, but how these aspects of the response are coordinated is poorly understood. We profiled the molecular and cellular features of the antibody response to influenza vaccination by integrating single-cell transcriptomics, longitudinal antibody repertoire sequencing, and antibody binding measurements. Single-cell transcriptional profiling revealed a program of memory B cell activation characterized by *CD11c* and *T-bet* expression associated with clonal expansion and differentiation toward effector function. Vaccination elicited an antibody clone which rapidly acquired broad high-affinity hemagglutinin binding during affinity maturation. Unexpectedly, many antibody clones elicited by vaccination do not bind vaccine, demonstrating non-specific activation of bystander antibodies by influenza vaccination. These results offer insight into how molecular recognition, transcriptional programs, and clonal proliferation are coordinated in the human B cell repertoire during memory recall.

## Main Text

Antibody memory is a hallmark of adaptive immunity and confers life-saving protection against many pathogens. During initial encounter with a pathogen, clonal selection and affinity maturation focus the antibody repertoire onto variants that bind specifically to pathogen-derived antigens with high affinity, and these antibodies are preserved in memory B cells. In subsequent encounters, memory B cells are rapidly activated, leading to clonal expansion and differentiation to antibody-secreting cells. This robust immune response can prevent reinfection or reduce severity of disease.

Although a protective memory response requires coordination of antigen recognition, gene expression, and clonal expansion, studies linking these facets of the response have been lacking. Specifically, deep sequencing-based measurements of the population dynamics and clonal structure of the B cell repertoire have shown that vaccination typically induces rapid expansion of a small set of B cell clones within 7 days (1–3). However, the transcriptional programs of these expanded clones and the antigen specificity of their antibodies have not been characterized.

Analogously, antigen-resolved measurements, such as serum binding assays and antigen-specific cell sorting, have demonstrated that antigen-specific serum antibody (4, 5), memory B cells (6), and antibody-secreting cells (7) become more abundant after vaccination. However, these approaches have not been able to resolve clonal relationships among antigen-specific cells, the population dynamics of these clones, or their gene expression programs.

Finally, bulk transcriptome measurements have detected transient expression signatures associated with memory recall after vaccination in blood (8, 9) and B cells (10), but it is not known how these transcriptional programs are related to clonal dynamics and antigen specificity within the B cell repertoire. Thus, an integrated portrait of how the memory response unfolds with cellular and molecular detail at the scale of the entire organism’s antibody repertoire remains lacking, despite its importance for protective immunity and vaccine design.

To address these questions, we developed an integrative approach that combines information from single-cell transcriptomics, longitudinal antibody repertoire sequencing, and antibody binding measurements, and applied it to study the human antibody response to influenza vaccination. We tracked the population dynamics of B cell clones in a time course after vaccination and profiled transcriptomes of single B cells within those clones, revealing an activated memory B cell state associated with vaccine-elicited clonal expansion. We then assessed the relationship between clonal expansion and antigen specificity by expressing native human antibodies isolated from single B cells and characterizing their binding properties.

### Integrating single B cell phenotypes with clonal population dynamics after vaccination

We studied the antibody repertoire response of one healthy young adult (age 18) to seasonal influenza vaccination in 2012. Deep multimodal study of a single individual’s vaccine response enabled us to extensively investigate the relationships between global repertoire structure and molecular function using a diverse suite of experimental techniques. To measure the population dynamics of the vaccine response, we sequenced the peripheral blood antibody repertoire (Rep-seq) at the time of vaccination and 1, 4, 7, 9, and 11 days afterward (D0, D1, D4, D7, D9, and D11), as well as 3 and 5 days before vaccination (D-3 and D-5) (Figure 1A and Figure 1B), as we previously reported (3). We detected ~625,000 unique antibody heavy chain sequences belonging to ~55,000 clonal lineages. Vaccination induced rapid recall of 16 vaccine-responsive clones, which were defined as those having >50-fold expansion in unique sequences detected between D0 and D7. These clones bear the hallmarks of memory B cells, including extensive somatic mutation, class-switched isotypes, and population genetic signatures of positive selection (3).

**Figure 1.**
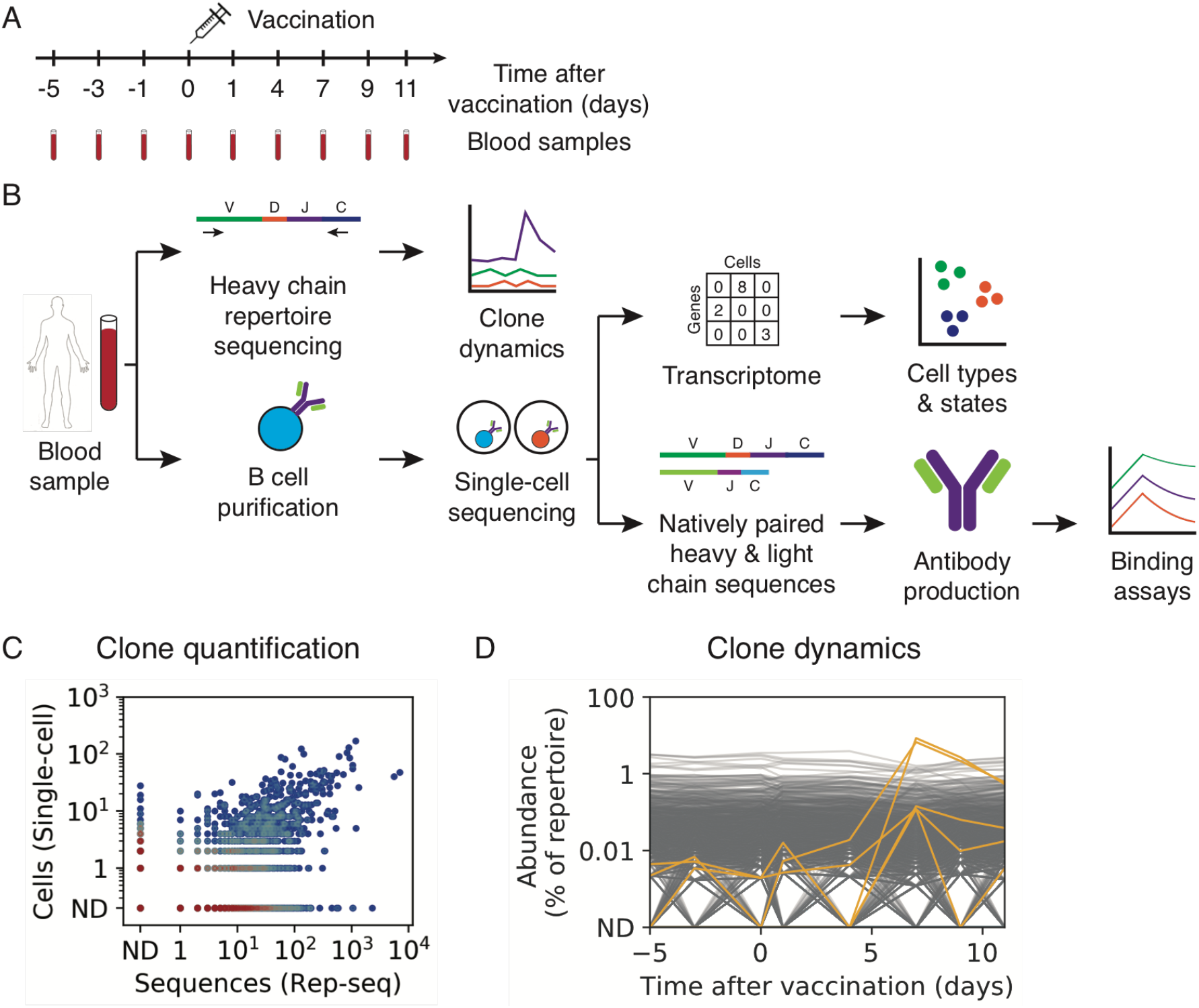
Characterization of B cell response to influenza vaccination using integrated single-cell and antibody repertoire sequencing. (**A**) Study design. (**B**) Experiment workflow. (**C**) Comparison of clonal abundance measurements across platforms, showing cells detected by single-cell sequencing and sequences detected by Rep-seq within each clone. Color indicates density of clones. ND, not detected. (**D**) Population dynamics of B cell clones. Each line shows a clone. Yellow lines indicate vaccine-responsive clones (>50-fold expansion from D0 to D7 and >0.1% of repertoire at D7).

We also sequenced antibody heavy and light chain transcripts in single B cells purified from peripheral blood samples of the same subject at D7 and D9, which correspond to the peak of the memory response (Figure 1B). After quality filtering and computational removal of doublets, we obtained 94,259 single B cells having exactly one productive heavy chain and one productive light chain transcript (Figure S1A and Figure S1B). We detected cells producing antibodies of every class, and the majority of cells produced IgM antibodies, as expected from pan-B cell purification, which includes naïve B cells (Figure S1C).

To connect single-cell phenotypes with clonal population dynamics, we mapped these single B cells to clones detected by Rep-seq using an established approach for identifying clonal lineages via single-linkage clustering (Figure S1D) (11, 12). Clones were identified in the Rep-seq repertoire for 8% of cells, with the nearest heavy chain complementarity determining region (HCDR3) exhibiting high identity (97% ± 3%, mean ± s.d.) for these matches (Figure S1E). Matches were strongly enriched for class-switched isotypes and depleted for IgD, as expected for memory B cells (Figure S1F). The majority of cells did not match a clone in Rep-seq data because most cells are naïve B cells, as confirmed by transcriptome profiling below. Additionally, the resampling probability of low abundance memory B cell clones across replicate samples is low (1). Nevertheless, for clones detected in both measurements, quantification of clone size was highly consistent across the two methods (Figure 1C; Spearman’s rho = 0.57, P < 10^−91^).

Based on the Rep-seq measurement of clonal population dynamics, we identified five vaccine-responsive clones that both expanded dramatically after vaccination (>50 fold-change from D0 to D7) and contained sequenced single cells. This included the two globally most abundant clones at the peak of recall at D7, and each clone comprised >0.1% of the repertoire at D7 (range 0.1 – 8%) (Figure 1D). Antibodies in these vaccine-responsive clones were mostly IgG (94%) and had extensive somatic hypermutation (mutation density 3.8% ± 1.4%, mean ± s.d.). These results establish that the combination of longitudinal Rep-seq and single-cell sequencing captures a rich portrait of B cell population dynamics at the scale of the whole organism, and links single-cell phenotypes such as paired heavy-light chain antibody sequences with clonal population dynamics.

Because single-cell sequencing preserves the native pairing between heavy and light chain sequences, we were able to assess the fidelity of the widely-used strategy of clone identification based on heavy chain sequence alone by using the light chain as an independent marker of clonal identity. Light chain genes were highly concordant within the vast majority of clones, as evidenced by the majority light chain gene representing a very high proportion of cells within each clone (Figure S1G; median = 100%, mean ± s.d. = 90% ± 18% for light chain constant region genes; similar results were found for light chain V and J genes). We observed that a minority of clones (16%) had substantial impurity based on the presence of cells containing a plurality of different light chain genes. We determined that these impure lineages were strongly enriched for short HCDR3 sequences (Figure S1H; P = 3.8 × 10^−91^, Mann-Whitney U test; median HCDR3 length 14 AA in impure lineages, 16 AA in pure lineages) and usage of the *IGHJ4* gene, which contributes a longer templated insert to the HCDR3 and thus tends to reduce sequence diversity (Figure S1H; P = 5.1 × 10^−225^, Fisher’s exact test; 64% *IGHJ4* usage in impure lineages, 28% in pure lineages). We conclude that the fidelity of clone identification based on clustering of heavy chain sequences is high for most clones. Clone assignment errors predominantly arise from low diversity compartments of the repertoire, and assignment can be improved by using light chain sequences when pairing information is available.

### Transcriptional program of vaccine-induced memory B cell activation

We performed single-cell transcriptome profiling on 35,631 cells, comprising a subset of the cells for which we sequenced antibody transcripts (Figure 2A). We detected a median of 2,015 UMIs and 766 genes per cell (Figure S2A), as typical for microfluidic droplet-based single-cell sequencing (13). Similar transcriptional profiles were obtained across 4 technical replicates (Figure S2B) and these data were pooled for analysis. Using t-SNE visualization and DBSCAN clustering, we identified distinct immune cell types, which we manually annotated based on established type-specific genes (Figure 2B). Three clusters corresponded to CD4+ and CD8+ T cells and macrophages, which displayed specific expression of markers such as *CD3E* for T cells and *LYS* for macrophages (Figure S2D) and lacked antibody expression (Figure 2C). These cell types were present at low abundance due to the imperfect purity of B cell isolation and were not analyzed further. B cells formed two distinct clusters, which we annotated as memory B cells and naive B cells based on established markers and antibody isotype (Figure 2B). Memory B cells expressed *CD27* (Figure S2E) and made predominantly class-switched antibodies (Figure 2C and Figure S2C). Naive B cells expressed *TCL1A* (Figure S2E) and made exclusively IgM and IgD antibodies (Figure 2C and Figure S2C). In total, we analyzed 16,653 memory and 18,953 naive B cells.

**Figure 2.**
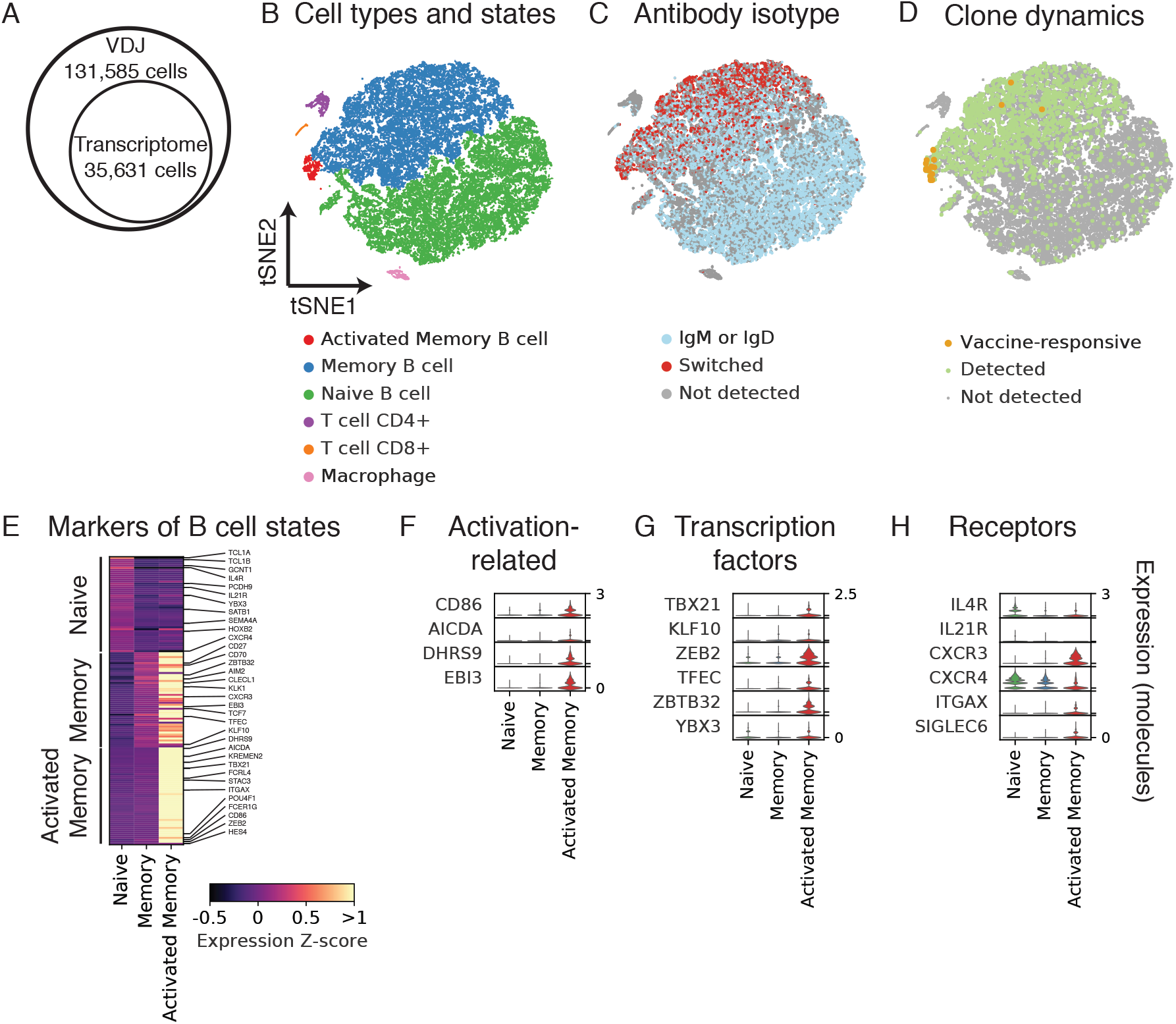
Characterization of gene expression in single B cells isolated from peripheral blood after influenza vaccination. (**A**) Number of cells analyzed using single-cell antibody gene sequencing (VDJ) or transcriptome profiling. (**B**–**D**) Principal components analysis and t-distributed Stochastic Neighbor Embedding (tSNE) separates cells into distinct clusters. Each dot is a cell, colored by type or state revealed by gene expression profile (**B**), antibody isotype as revealed by antibody sequencing (**C**), or clonal population dynamics as revealed by Rep-seq (**D**). (**E**) Differential expression analysis identifies markers of distinct B cell states. Genes of immunological interest are labeled. (**F**–**H**) Gene expression distributions in distinct B cell states of established immune activation-related genes (**F**), transcription factors (**G**), and signaling receptors (**H**).

To address how clonal population dynamics are related to transcriptome state, we mapped the single B cell transcriptomes to the clones identified using Rep-seq based on heavy chain sequence, as described above. Matches to clonal lineages were obtained almost exclusively for memory B cells as expected (Figure 2D). Remarkably, we found that cells belonging to vaccine-responsive clones had a distinct transcriptional profile characteristic of a small neighborhood within the memory B cell cluster (Figure 2D). Cells in this neighborhood expressed established genes related to B cell activation, including the activation marker *CD86* and the somatic hypermutation gene *AICDA*, also known as *AID* (Figure 2E, Figure 2F, and Figure S2E). Thus we annotated cells in this neighborhood as activated memory B cells, comprising 421 cells in total.

To define the transcriptional programs of B cell states, we identified genes exhibiting differential expression across naive, memory, and activated memory B cells. We found 755 differentially expressed genes between naive and memory B cells (FDR = 0.1%, Mann-Whitney U test with Benjamini-Hochberg correction), including established markers such as *CD27* and *IGHD* (Figure 2E). About half of these genes were upregulated in naive B cells, while the other half were upregulated in memory B cells (Figure S2G). By contrast, we found 172 differentially expressed genes between memory and activated memory B cells, all of which were upregulated in activated memory B cells (Figure S2G). Dominant upregulation of genes in the activated memory state was consistently observed across a range of significance thresholds defining differential expression (Figure S2G). We also detected more genes (median 1,786) and more UMIs (median 5,517) in activated memory B cells than memory B cells (Figure S2F; median gene count in memory B cells = 849, UMI count = 2,406), possibly reflecting greater mRNA content due to elevated transcription. Together, these results suggest that the transcriptional program of memory B cell activation predominantly involves activation rather than deactivation of gene expression.

To characterize the activated memory B cell state, we first sought to identify transcription factors (TFs), which may be central regulators of the program of activation. We identified 6 TFs specifically expressed in activated memory B cells (Figure 2G). These TFs include *T-bet*, also known as *TBX21* (Figure S2E), which is required for IgG2a class switching (14) and clearing chronic viral infections (15), and *Zbtb32*, which modulates the duration of memory B cell recall responses in mice (16).

Several cytokine receptors are downregulated in activated memory B cells (Figure 2H). *IL4R* and *IL21R* are highly expressed in naive B cells, but downregulated in memory and activated memory B cells (Figure 2H and Figure S2E), suggesting that naive B cells are more responsive than memory or activated memory B cells to IL4 and IL21, which regulate class switching to IgG4 or IgE (17), and IgG1 or IgG3 (18), respectively. The chemokine receptor *CXCR4*, which controls entry to anatomical locations of B cell maturation, such as lymph nodes and Peyer’s patches (19), is also progressively downregulated from naive to memory and activated memory B cells (Figure 2H).

Other genes related to humoral activation are upregulated in activated memory B cells. The chemokine receptor *CXCR3*, which is required for cell migration to sites of inflammation (20), is specifically expressed in activated memory B cells (Figure 2H and Figure S2E). Interestingly, *CD11c*, also known as *ITGAX*, is specifically expressed in activated memory B cells (Figure 2H), suggesting that this state overlaps with the recently described age/autoimmune-associated B cells (21, 22). Finally, *EBI3*, which is known to be expressed in germinal center B cells (23), is found exclusively in activated memory B cells (Figure 2F). Complete lists of differentially expressed genes across naive, memory, and activated memory B cells are shown in Table S1 and Table S2. Together, these results define a transcriptional program of memory B cell activation associated with vaccine-induced clonal expansion, which bears hallmarks of an effector B cell response.

### Many vaccine-responsive antibodies do not bind vaccine

To study how clone dynamics and antigen specificity are related, we expressed and functionally characterized 21 antibodies obtained from single B cells within 5 vaccine-responsive clones (Figure S3A). We first measured binding of these antibodies to the vaccine (trivalent influenza vaccine from the 2011–2012 flu season) by ELISA. Surprisingly, only 57% of the vaccine-responsive antibodies (12 of 21) and 40% of vaccine-responsive antibody clones (2 of 5) exhibited binding to vaccine (Figure 3 and Figure S3B). For the non-vaccine-binding antibodies, we further screened for binding by ELISA against a panel of purified influenza proteins, including hemagglutinins, neuraminidases, nucleoprotein, matrix protein, and non-structural proteins, but found no binding (Figure S3C). Notably, despite not binding vaccine or influenza proteins, these three clones expanded dramatically after vaccination (>62-fold) and were highly abundant at D7, including one clone which was the second most globally abundant clone, representing 6.7% of the repertoire at D7. These results indicate that many vaccine-responsive antibodies do not bind vaccine or purified components of the vaccine. This suggests that vaccination induced activation of some antibody clones in an antigen-independent manner. We found no binding of these non-vaccine-binding antibodies to a panel of common viral and bacterial antigens, such as herpes simplex, measles, and varicella zoster virus (Figure S3C), and we were unable to determine the specificities of these antibodies.

**Figure 3.**
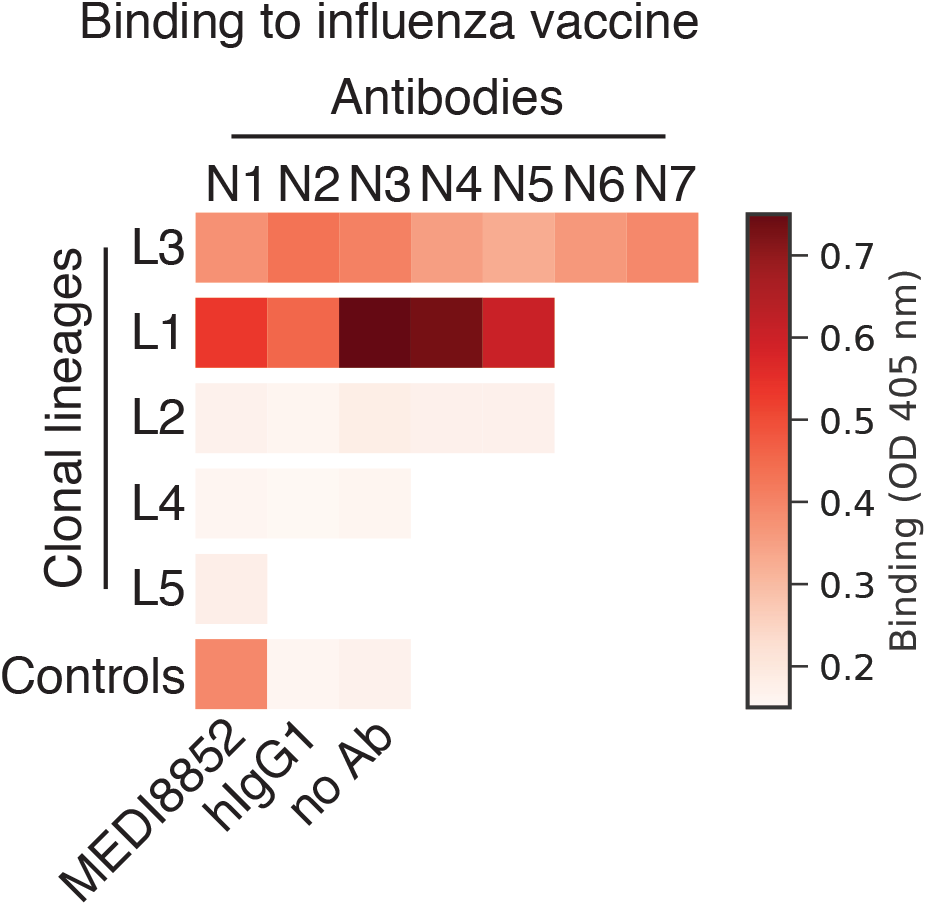
Binding of influenza vaccine-responsive antibodies to vaccine. Binding of 21 monoclonal antibodies from 5 clones to the trivalent inactivated influenza vaccine from 2011–2012 season was measured using enzyme-linked immunosorbent assay (ELISA), revealing that many vaccine-responsive antibodies do not bind vaccine. OD, optical density; hIgG1, human IgG1.

### A broadly binding high-affinity anti-influenza antibody clone elicited by vaccination

To determine the specificity of the vaccine-binding antibodies, we screened them for binding by ELISA against purified influenza proteins, including the major antigenic determinants of influenza virus, hemagglutinin (HA) and neuraminidase (NA). One vaccine-responsive clone, which we refer to as L3, displayed strong binding to diverse HA proteins, including the influenza A variants contained in the vaccine, H1 A/California/7/2009 and H3 A/Perth/16/2009, as well as H5 and H9 variants (Figure 4A). These antibodies had similar binding strength and breadth as established broadly neutralizing antibodies MEDI8852 (24) and CR9114 (25) (Figure 4A). L3 antibodies use the *IGHV3-34* and *IGHJ4* genes, have a 19 AA HCDR3, and are heavily mutated (28 ± 5 mutations from inferred germline heavy chain, mean ± s.d.) (Figure S3A).

**Figure 4.**
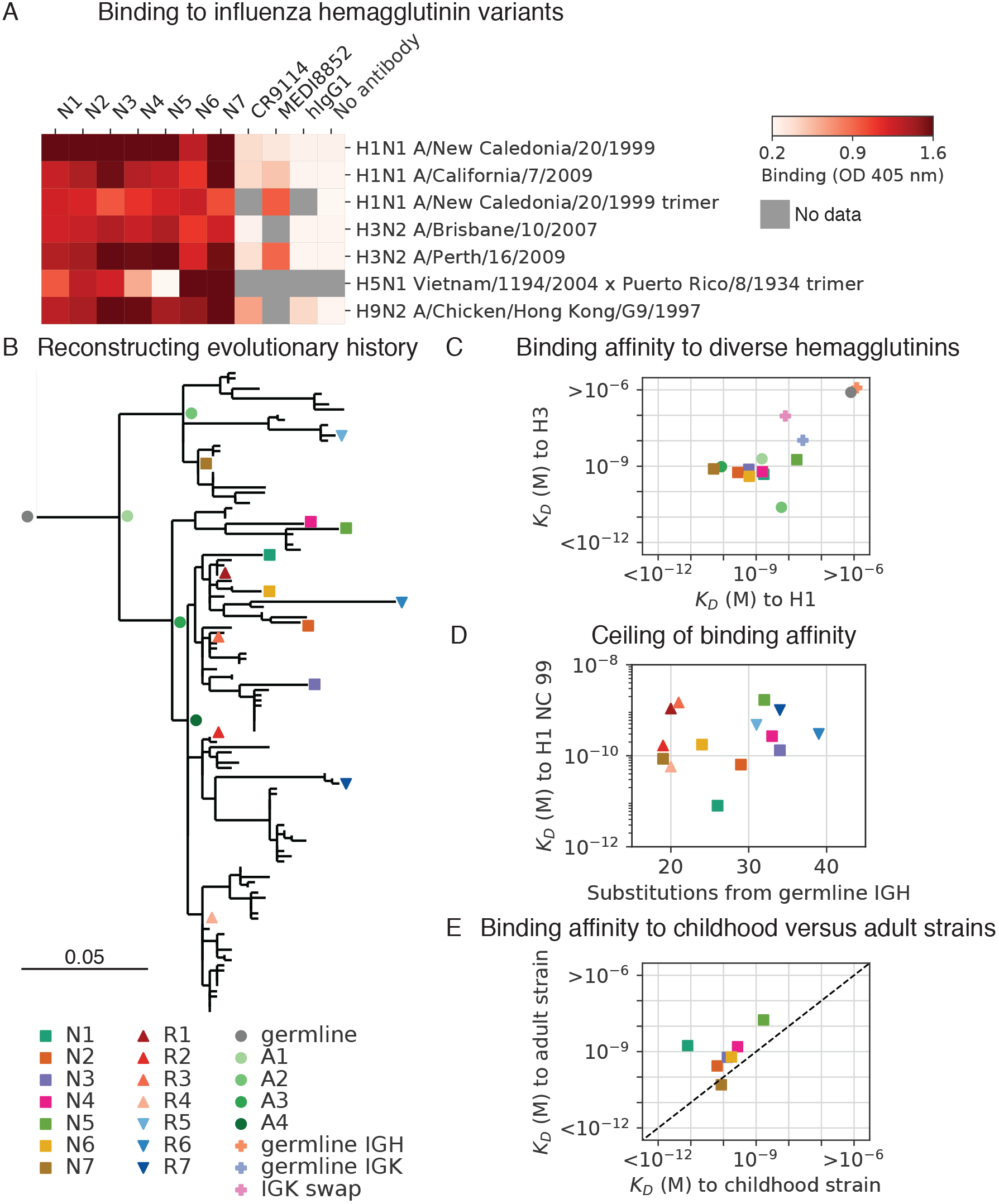
Reconstructing evolution of a broadly binding high-affinity anti-influenza antibody clone. (**A**) Binding of antibodies from the L3 clone to a panel of influenza hemagglutinin (HA) variants was measured using ELISA. OD, optical density; hIgG1, human IgG1. (**B**) Evolutionary history of L3 depicted as a maximum-likelihood phylogeny based on heavy chain sequence. Markers indicate antibodies detected by single-cell sequencing (N1–7) or repertoire sequencing (R1–7), or reconstructed ancestral sequences (germline and A1–4). (**C**) Equilibrium constants (K_D_) of binding between L3 antibody variants and H1 (A/California/7/2009) and H3 (A/Perth/16/2009) hemagglutinin variants, as determined by biolayer interferometry. L3 antibodies include extant sequences (N1–7), reconstructed ancestral sequences (germline and A1–4), and engineered variants having the L3N6 sequences, but with heavy chain reverted to the inferred germline sequence (germline IGH), light chain reverted to the inferred germline sequence (germline IGK), or a light chain sequence substituted from a different clone (IGK swap). Jitter was added to germline and germline IGH to improve visualization of the data points. (**D**) Equilibrium constants of binding between L3 antibodies compared with extent of somatic hypermutation. (**E**) Equilibrium constants of binding between L3 antibodies and H1 variants from childhood (A/New Caledonia/20/1999) and adulthood (A/California/7/2009). Dashed line indicates equal K_D_ for binding to both variants.

We measured the binding affinity of L3 antibodies to diverse H1 and H3 variants using biolayer interferometry. Most L3 antibodies bound with sub-nanomolar affinity to both H1 and H3, which are highly divergent HA variants drawn from the two major groups of influenza A virus and share only 44% amino acid identity (Figure 4C and Figure S4C; equilibrium binding constants [K_D_] from 18 nM to 50 pM). Thus, L3 broadly binds diverse hemagglutinin variants with high affinity. A second vaccine-binding clone, which we call L1, displayed strong but narrow binding specificity to HA B (Figure S3C) and we did not analyze this clone further.

### Evolution of a broadly binding anti-influenza antibody clone

To shed light on the evolutionary trajectories leading to broad high-affinity anti-influenza binding, we reconstructed the clonal evolution of L3 (Figure 4B). Using maximum-likelihood phylogenetic models, we reconstructed the ancestral sequences of the unmutated germline precursor and four intermediate ancestors (Figure S4A and Figure S4B), then expressed these antibodies and measured their binding affinities to diverse HAs. While the germline precursor bound weakly to H1 and H3 (K_D_ > 1 uM) (Figure S4E), the first intermediate ancestor A1 bound to both H1 and H3 with nanomolar affinity (K_D_ = 1.5 nM and 2 nM, respectively) (Figure 4C), despite having acquired only 11 amino acid substitutions (6 in the heavy chain and 5 in the light chain) (Figure S4A).

To dissect the contributions of heavy and light chain mutations to binding affinity, we engineered variants of the high-affinity L3N6 antibody in which the heavy and light chain sequences were separately reverted to the respective germline precursor sequence (Figure S4A and Figure S4B). We found that germline reversion of the heavy chain greatly reduced binding affinity to both H1 and H3 (K_D_ > 1 uM) (Figure 4C and Figure S4E). In contrast, germline reversion of the light chain minimally affected binding to H1 and H3 (K_D_ = 27 nM and 10 nM, respectively) (Figure 4C and Figure S4E). To further test the contribution of light chain mutations, we created a variant of L3N6 in which the light chain was swapped for a different IGK sequence originating from a distinct clonal lineage having a different LCDR3 (Figure S4B). This alteration of the light chain also minimally affected binding to H1 and H3 (K_D_ = 8 nM and 94 nM, respectively) (Figure 4C and Figure S4E). These findings show that heavy chain mutations were predominantly responsible for affinity maturation, indicating that broad nanomolar-affinity binding was achieved via ≤6 amino acid substitutions in the heavy chain.

L3 antibodies therefore rapidly evolved broad high-affinity binding to diverse HA variants through a small number of somatic mutations. Affinity improvements were predominantly driven by decreasing the dissociation rate, which varied ~10,000-fold across the clone, rather than increasing the association rate, which varied only ~10-fold (Figure S4D). We found evidence for an affinity ceiling: acquisition of mutations beyond the intermediate ancestor A1 did not substantially affect affinity and there was no trend toward enhanced affinity with additional mutations (across the range of 18 – 38 mutations from the inferred germline IGH sequence) (Figure 4D; Spearman’s rho = 0.25, P = 0.37). Instead, L3 antibody affinity evidently drifted neutrally after acquisition of high-affinity binding.

To determine how L3 antibodies bind HA, we performed cross-competition binding experiments using biolayer interferometry. We competed L3N1 and L3N6 against a panel of broadly binding antibodies consisting of stem-binding antibodies CR9114 (25) and MEDI8852 (24), receptor-binding site antibodies CH65 (26) and H2897 (27), and lateral patch antibody 6649 (28). We found that L3N1 and L3N6 did not compete with any of these antibodies (Figure S5). This result indicates that the epitopes recognized by L3 antibodies do not overlap with any antibodies in this panel, suggesting that L3 achieves broad specificity by a distinct structural mechanism. Furthermore, the L3 epitope may be conserved across HA variants belonging to groups 1 and 2.

It has been proposed that the antibody memory response is biased towards antigens seen early in an individual’s life, and this priming influences subsequent responses (29). To test this hypothesis using L3, we compared binding affinity to H1 variants that circulated during the subject’s childhood and adulthood. We found that extant antibodies of the L3 clone nearly all bound with higher affinity to the childhood strain (H1 New Caledonia/20/1999) than the adult strain (H1 California/07/2009) (Figure 4D). This indicates that the affinity of a broad binding anti-HA antibody clone is biased towards antigenic variants associated with childhood exposure, supporting the hypothesis that affinity maturation most efficiently focuses the antibody repertoire on antigens encountered in early life, leaving an lasting imprint on subsequent responses.

## Discussion

Mobilization of an effective antibody memory response requires coordination across scales, from antibody-antigen recognition and transcriptional activation in single cells to clonal population dynamics that globally remodel an organism’s antibody repertoire. This complex, multi-scale nature of the immune system creates challenges for understanding its function. To address these challenges, we have developed an experimental approach that integrates single B cell sequencing with longitudinal antibody repertoire sequencing and biophysical measurements of antibody function. Our results show that this strategy offers a unified portrait of the molecular and cellular features of the memory B cell response to vaccination, giving insights into mechanisms of immune memory.

Much recent interest has focused on a functionally specialized B cell subset marked by *CD11c* and *T-bet* expression named “Age/autoimmune Associated B cells” (ABCs). B cells with these features are associated with viral infections, autoimmunity, and aging in mouse and human (21, 22, 30–32), but to our knowledge the phenotype has not been described as a transcriptional state with single-cell resolution. Using single-cell transcriptomics and longitudinal clone tracking, we have defined an activated memory B cell state, which displays hallmarks of an effector B cell response and shares many features with ABCs, including high expression of *CD11c* (21, 22), *T-bet* (31), *FCRL4*, and *CXCR3* (30). Several genes that define this activated memory B cell state are directly involved in germinal center migration (*EBI3*) (23), somatic hypermutation (*AICDA*) (33), and class switching (*AICDA* and *Tbx21*) (14, 33), suggesting that these cells are poised for secondary affinity maturation. Our results indicate that these *CD11c*+ *T-bet*+ B cells are associated with vaccine-elicited clonal expansion in a healthy young adult human. These findings support the view that *CD11c*+ *T-bet*+ B cells are essential to health, but aberrant regulation of them can lead to autoimmunity. Defining their transcriptional program opens avenues to understanding their origins, function, and regulation, which may in turn reveal therapeutic targets in both pathogen immunity and autoimmunity.

Unexpectedly, several antibody clones elicited by vaccination did not bind vaccine. Formally, we cannot exclude that the lack of binding between recombinant vaccine-responsive antibodies and the vaccine in vitro is due to conformational changes occurring under physiological conditions. Notwithstanding this alternative explanation, our results suggest that bystander activation of memory B cells bearing non-vaccine specificities is common after influenza vaccination. Polyclonal activation of memory B cells bearing non-vaccine specificities after vaccination has previously been described at the level of serum antibody (34) and antibody-secreting cells (7). Similarly, infection with both measles and varicella induces non-specific B cell activation (35). Our results show that many, perhaps even the majority of, memory B cells elicited by influenza vaccination produce antibodies that do not bind the vaccine, revealing an unanticipated extent of this phenomenon. This extent comports with some previous studies based on single-cell cloning of antibody-secreting cells (7), but may have been underestimated in other studies that tested binding against limited panels of antigens (34, 36). Non-specific polyclonal activation has been proposed as a mechanism for maintenance of long-term immune memory, enabling memory cell proliferation in the absence of antigen encounter (37). We were not able to identify antigens for the non-vaccine-binding antibodies by screening against a panel of common viral and bacterial antigens; conclusive identification of non-vaccine specificities will require high-throughput screening methods. Nevertheless, our integrated strategy of single-cell and Rep-seq offers a direct route to characterization of these non-vaccine-specific yet vaccine-elicited antibodies. Our results also indicate that bystander activation is confined to a small number of clones by an unknown mechanism, perhaps related to the presence of activated T cells (38–40).

We discovered a broadly binding anti-hemagglutinin antibody clone in which fewer than six somatic mutations in the heavy chain alone was sufficient to confer broad high-affinity binding, offering a striking example of rapid affinity maturation. Together with prior examples of influenza antibodies that emerged via a small number of mutations (41, 42), this suggests that a single-dose vaccine could be sufficient to confer lasting protection to influenza. Unlike prior examples (41, 42), L3 does not use the heavy chain variable region *VH1-69* gene, potentially opening a new target for germline-targeting immunogens. L3 antibodies may bind a distinct epitope compared with previously identified classes of broadly binding anti-HA antibodies (24–28), suggesting that structural characterization of the interaction may reveal a new site of vulnerability on HA.

## Acknowledgments

We thank Peter Kim and Derek Croote for helpful discussions, 10X Genomics for providing reagents for single-cell sequencing, Krista McCutcheon for expressing antibodies, Payton Weidenbacher for discussions and providing antibodies and influenza proteins, and Harry Greenberg and Caiqiu Zhang for providing trivalent inactivated influenza vaccine. This work was supported by NIH U19A1057229 (S.R.Q.) and the National Science Foundation Graduate Research Fellowship Program (F.H.). The authors declare no competing interests.

## Supplementary Materials

## Materials and Methods

### Study subject

Study subject gave informed consent and protocols were approved by the Stanford Institutional Review Board. Subject was a female human aged 18 who was recruited in 2012. The subject was apparently healthy and showed no signs of disease.

### Sample collection

As previously described (3), blood was drawn by venipuncture, then peripheral blood mononuclear cells (PBMCs) were isolated using a Ficoll gradient and frozen in 10% (vol/vol) DMSO and 40% fetal bovine serum (FBS) according to Stanford Human Immune Monitoring Center protocol. Subject was vaccinated with the 2011–2012 seasonal trivalent inactivated influenza vaccine. Blood was collected 3 and 5 days before vaccination (D-3 and D-5); immediately before vaccination (D0); and 1, 4, 7, 9, and 11 days afterwards (D1, D4, D7, D9, D11).

### Antibody repertoire sequencing

Antibody repertoire sequencing was previously performed on samples from all timepoints and preprocessed data was downloaded (3). Briefly, PBMCs were thawed and RNA was extracted. This RNA was reverse transcribed using immunoglobulin heavy chain constant region-specific primers and cDNA was amplified by PCR. UMIs were incorporated during reverse transcription and PCR. These libraries were sequenced using the Illumina HiSeq 2500 and MiSeq platforms using paired-end 101 or 300 bp reads, respectively. Consensus-based error correction was performed using UMIs. Sequences were annotated with V and J germline gene usage using IgBlast (43) and isotype using BLASTN (44). Clonal lineages were identified based on V and J gene usage, HCDR3 length, and HCDR3 sequence composition. Dynamics of clones were determined by comparing fractional abundance across study timepoints. As in our previous study (3), vaccine-responsive clones were identified as those having >50-fold expansion from D0 to D7 and composing >0.1% of the repertoire at D7.

### Single-cell isolation and sequencing

PBMCs from D7 and D9, which correspond to the peak of the B cell memory recall response, were thawed. B cells were magnetically enriched using the B Cell Isolation Kit II (Miltenyi). Single cells were encapsulated in droplets using 16 lanes of the Chromium device (10X Genomics) with target loading of 14,000 cells per lane. Reverse transcription and cDNA amplification were performed using the Single Cell V(D)J kit (10X Genomics). In 12 lanes, direct enrichment of VDJ was performed. In the remaining 4 lanes, VDJ and gene expression measurement was performed; these 4 lanes were considered technical replicates. All steps were done according to manufacturer’s instructions, except with additional cycles of polymerase chain reaction (PCR) (19 total cycles for direct enrichment of VDJ; 22 total cycles for VDJ and gene expression). 50 ng of cDNA was used as input for library preparation. Libraries were sequenced using the Illumina NextSeq 500 platform with paired-end reads for VDJ of 150 bp each and for gene expression of 26 bp and 98 bp.

### Preprocessing of single-cell sequence data

Sequences were preprocessed to map reads to the human reference genome (GRCh38) using STAR 2.5.1b (45), count molecules aligning to each gene, and assemble antibody heavy and light chain transcripts within cellranger 2.1.0. To distinguish bona fide single cells from multiplets, we examined the number of productive heavy and light chain contigs assembled for each cell barcode. Single B cells were identified by the presence of a single productive heavy chain and a single productive light chain, yielding a total of 94,259 single B cells for analysis. All other cells were excluded from further analysis.

### Mapping single B cells into clones

Single B cells were mapped to clones using a custom algorithm similar to that used for identification of clones previously (3, 11). Sequences detected by repertoire sequencing (n = 625,750) were annotated for V and J gene usage, HCDR3 length, and HCDR3 sequence and formed the database of subject sequences. The heavy chain variable region sequence from each single B cell was used as a query to search this database. For each query, the set of subjects sharing the query’s V and J genes and CDR3 length was identified. Within this set, the identity between the query and subject sequences within the HCDR3 and outside the HCDR3 were calculated based on Hamming distance, and hits were defined as having >90% nucleotide identity in both regions. Previous studies have demonstrated that this cutoff of sequence identity enables identification of clonally related sequences with high sensitivity and specificity (11, 12). This yielded 8,377 single B cells that had matching clones detected by repertoire sequencing.

Fidelity of clonal clustering was assessed using the light chain as an independent marker of clonal identity. In clones having multiple B cells detected by single-cell sequencing, the percentage of cells possessing the dominant light chain was determined. Impure clones were identified as those having <80% of cells within the clone sharing the dominant light chain. All of the vaccine-responsive clones were pure.

### Analysis of gene expression in single cells

Gene expression profiles were log-transformed and normalized to counts per million molecules. Dimensionality reduction using principal components analysis (PCA) retaining the top 10 principal components followed by t-distributed Stochastic Neighbor Embedding (tSNE; perplexity = 30, theta = 0.5, max_iter = 1,000) (46) were performed using cellranger 2.1.0. Clusters were identified using Density-Based Spatial Clustering of Applications with Noise (DBSCAN; eps = 0.66, min_samples = 10) (47) and annotated based on expression of established marker genes for each cell type. Differentially expressed genes were identified using the negative binomial exact test adjusted for multiple testing using the Benjamini-Hochberg procedure as implemented in Loupe 2.0.0 (10X Genomics). For visualization of differential expression, the Z-score of expression of each group of cells was computed in comparison with the mean and standard deviation of expression in all other cells. Data visualization and analysis were performed using Scanpy (48) within JupyterLab (49).

### Reconstructing the evolutionary history of antibody clone L3

Evolutionary analysis was conducted sequences in clone L3 obtained by repertoire sequencing using paired-end 300 bp reads (n = 125) and single-cell sequencing (n = 7). Sequences were initially aligned in an ungapped manner using the start and end positions of the HCDR3 as anchor points. This alignment was refined using MUSCLE 3.8.31 with “-refine -maxiters 1 -diags -gapopen −5000” (50), then trimmed to remove positions which were only covered by single-cell sequencing contigs (which are substantially longer than repertoire sequencing assemblies). We added an inferred germline sequence consisting of the reference heavy chain V and J genes and the consensus of the alignment for the untemplated regions of the HCDR3. Phylogenetic reconstruction was performed using FastTree 2.1.7 with “-nt -gtr” (51). We concatenated light chain sequences to this alignment, then performed reconstruction by maximum-likelihood assuming equal rates for all mutations.

To assess the contribution of heavy and light chain mutations to binding, we engineered variants of the high-affinity antibody L3N6 by substituting either the inferred germline heavy (germline IGH) or light (germline IGK) chain sequence. We also substituted the light chain with a randomly chosen sequence from a different clonal lineage that used the same VK gene, but had a distinct LCDR3 (IGK swap). For cloning and expression of antibodies derived from repertoire sequencing (R1–7), we used the light chain sequence originating from the single cell nearest the selected antibody, using the metric of heavy chain nucleotide sequence identity. These antibodies were chosen to span a wide range of somatic mutation levels.

### Recombinant antibody expression

Recombinant antibodies were cloned and expressed by Genscript. Briefly, selected antibodies were codon-optimized for human expression. These DNA sequences were synthesized and cloned into heavy and light chain pcDNA3.4 expression vectors. Heavy chains were expressed as human IgG1 and light chains were expressed as either human IGK or IGL as appropriate. Vectors were transiently transfected in HEK293-6E cells and antibodies were purified from supernatant using Robocolumn Eshmuno A (EMD Millipore) or Monofinity A Resin prepacked columns (Genscript). Purity generally >95% was confirmed using SDS-PAGE and immunoblots under reducing and non-reducing conditions.

### Antigens for binding measurements

Fluzone trivalent inactivated influenza vaccine from the 2011–2012 flu season (Sanofi Pasteur) containing H1N1 A/California/7/2009, H3N2 A/Perth/16/2009, and B/Brisbane/60/2008 was obtained as a gift from Dr. Harry Greenberg. Purified influenza proteins expressed in human cells (typically HEK293) where possible, otherwise baculovirus or E. coli, were purchased from Sino Biological (11683-V08H, 11085-V08H, 11056-V08H, 40043-V08H, 11048-V08H, 40104-V08H, 40036-V08H, 11053-V08H, 40197-V07H, 40017-V07H, 40569-V07H, 40502-V07B, 40205-V08B, 40499-V08B, 40010-V07E, 40107-V08E, 40011-V07E, 40012-VNAE). Viruses inactivated by irradiation or formaldehyde treatment were purchased from Biorad (PIP005, PIP009, PIP010, PIP013, PIP014, PIP023, PIP008, PIP015, PIP016). Tetanus toxin was purchased from Sigma Aldrich (T3194).

### Binding measurements using ELISA

Semi-quantitative measurements of binding were carried out using enzyme-linked immunosorbent assay (ELISA). Antigen was immobilized on clear polystyrene 96- or 384-well MaxiSorp plates (ThermoFisher) by overnight incubation at 4 C at 2 ng/uL diluted in phosphate-buffered saline (PBS) pH 7.4, then three washes were performed. When vaccine was used as antigen, vaccine was immobilized at a 50-fold dilution in PBS pH 7.4. The plate was incubated for 2 hours at room temperature with blocking buffer (PBS pH 7.4 with 0.05% Tween-20 and 2% bovine serum albumin [BSA]), then washed twice. The plate was incubated with primary antibody diluted to 2 ng/uL unless otherwise noted in blocking buffer for 2 hours at room temperature, then washed four times. The plate was incubated with detection antibody (mouse anti-human IgG1 Fc conjugated to horseradish peroxidase clone HP6069; ThermoFisher) for 2 hours at room temperature, then washed five times. All washes consisted of 5 minute incubation with PBS pH 7.4 with 0.05% Tween-20. Detection was performed by adding 1-Step ABTS Substrate (ThermoFisher), then measuring absorbance at 405 nm at 1 or 3 min intervals for 45 min. Time point used for analysis was determined based on the dynamic range of the data (increasing signal, but no saturation). Positive controls included the broadly binding anti-influenza antibodies MEDI8852 (24) and CR9114 (25) obtained as a gift from Dr. Peter Kim. As negative controls, we used natural human IgG1 prepared from myeloma plasma (Abcam), or incubated wells with PBS alone instead of antigen (referred to as “no antigen”) or blocking buffer alone instead of antibody (referred to as “no antibody”).

### Binding measurements using biolayer interferometry

Kinetic measurements of antibody-antigen interactions were performed using biolayer interferometry on a ForteBio Octet 96 instrument with anti-human IgG Fc capture (AHC) biosensors. All assays were carried out in PBS with 1% BSA and 0.05% Tween-20 with a total volume of 250 uL per well using the following protocol: 60 s baseline, 300 s loading of antibody, 60 s baseline, 300 s association of antigen, and dissociation of variable duration up to 600 s for high affinity interactions. Antibody was loaded at 1.5 ng/uL and antigen concentrations ranged from 2.5 to 100 nM. Between assays, sensors were regenerated by cycling between assay buffer and 10 mM glycine pH 1.5 for 30 s, then quenched for 30 s in assay buffer. Data were processed using ForteBio software and custom Python scripts to perform global fitting of a 1:1 binding model across 2–5 antigen concentrations after double reference subtraction (using buffer only and analyte only conditions).

To determine whether antibodies bind similar or overlapping epitopes, competitive binding of antibody pairs to a specific antigen was characterized using anti-penta-HIS (HIS1K) biosensors. We used the following protocol: 60 s baseline, 300 s loading of antigen, 60 s baseline, 900 s association of blocking antibody, 60 s baseline, 600 s association of test antibody. Antigen was HA H1N1 A/New Caledonia/20/1999 with an isoleucine zipper trimerization domain and polyhistidine tag obtained as a gift from Dr. Peter Kim and used at 25 nM. Blocking antibodies were used at 400 nM and included MEDI8852 (24), CR9114 (25), CH65 (26), H2897 (27), and 6649 (28) obtained as gifts from Dr. Peter Kim. Test antibodies were used at 100 nM and included L3N1 and L3N6. As a control, self-blocking assays were performed using the same antibody for blocking and test steps, except with test antibody at 100 nM. Data were processed using ForteBio software and custom Python scripts. We note that complete blocking was observed between MEDI8852 and CR9114, which have overlapping epitopes. Partial blocking was observed between 6649 and H2897, which have partially overlapping epitopes.

### Data and code availability

Sequence data, preprocessed data, and code will be made freely available at the time of publication.

---

**Figure S1.**
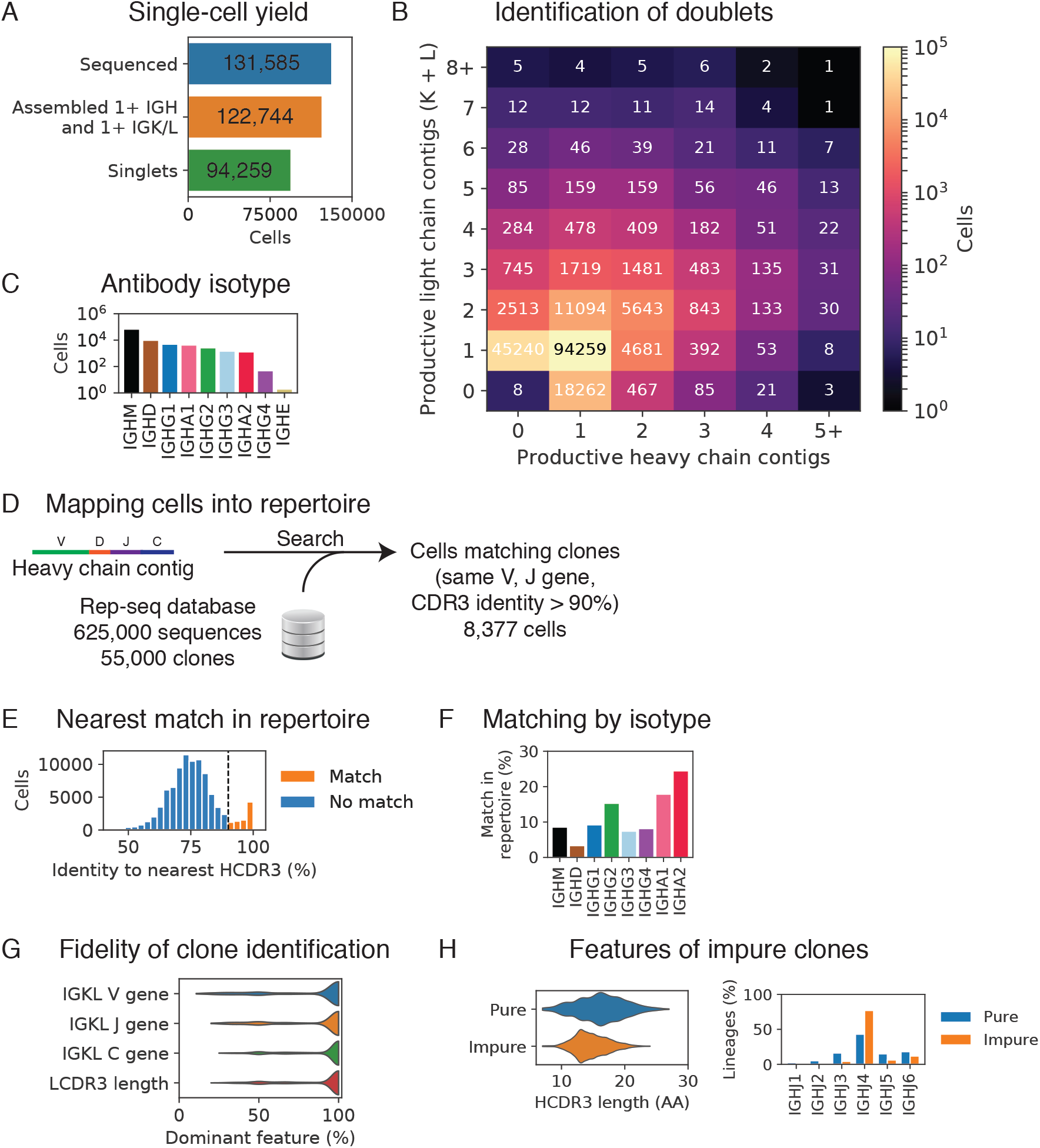
Performance and features of integrated single-cell and antibody repertoire sequencing measurements. (**A**) Number of single cells that were sequenced, had at least one productive heavy (IGH) and one light (IGK/L) chain gene assembled, or had exactly one productive heavy and one productive light chain gene assembled (singlets). (**B**) Doublets were identified and removed based on the number of productive heavy and light chain contigs assembled. (**C**) Isotype of antibodies from single B cells were determined based on the heavy chain constant region sequence. (**D**) Single B cells were mapped to clones detected by repertoire sequencing using a custom search algorithm. Matches required usage of the same V and J genes and HCDR3 identity >90%. (**E**) HCDR3 identity of nearest match in repertoire sequences. Dashed line indicates cutoff of 90% HCDR3 identity. (**F**) Isotypes of antibodies in the single cells that matched clones detected by repertoire sequencing (n = 8,377) (**G**) Fidelity of clone identification was determined by assessing the fraction of single cells sharing the characteristics of the dominant light chain gene found within the clone. (**H**) Molecular features of pure and impure clones (as determined based on light chain characteristics). AA, amino acids.

**Figure S2.**
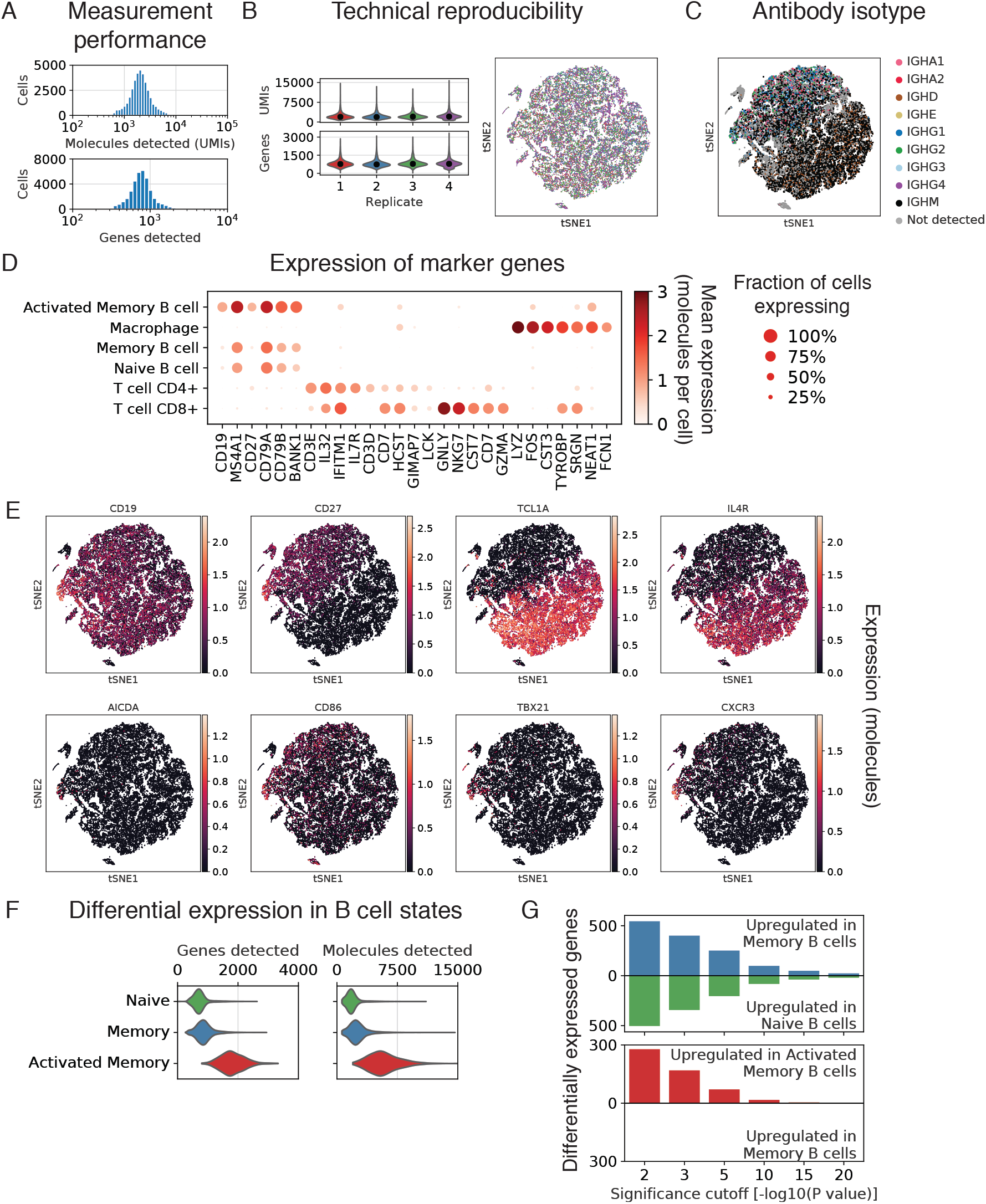
Additional characterization of gene expression in single cells isolated from peripheral blood after influenza vaccination. (**A**) Genes and molecules detected in each individual cell. UMIs, unique molecular identifiers. (**B**) Distributions of genes and molecules detected in individual cells in technical replicates. Median is indicated by black dot. In right plot, each dot is a cell colored by technical replicate of origin, according to the colors in left plots. tSNE, t-distributed Stochastic Neighbor Embedding. (**C**) Isotypes of antibodies in single cells were determined based on the heavy chain constant region gene. (**D**) Clusters were annotated as distinct cell types and states based on expression of established marker genes. Each dot is a cell. (**E**) Expression of selected established marker genes and genes of immunological interest in single cells. Each dot is a cell. (**F**) Distributions of genes and molecules detected in B cells in distinct states. (**G**) Differential expression analysis identified genes upregulated in naïve compared to memory B cells (green), memory compared naïve B cells (blue), and activated memory compared to memory B cells (red) across a range of significance cutoffs.

**Figure S3.**
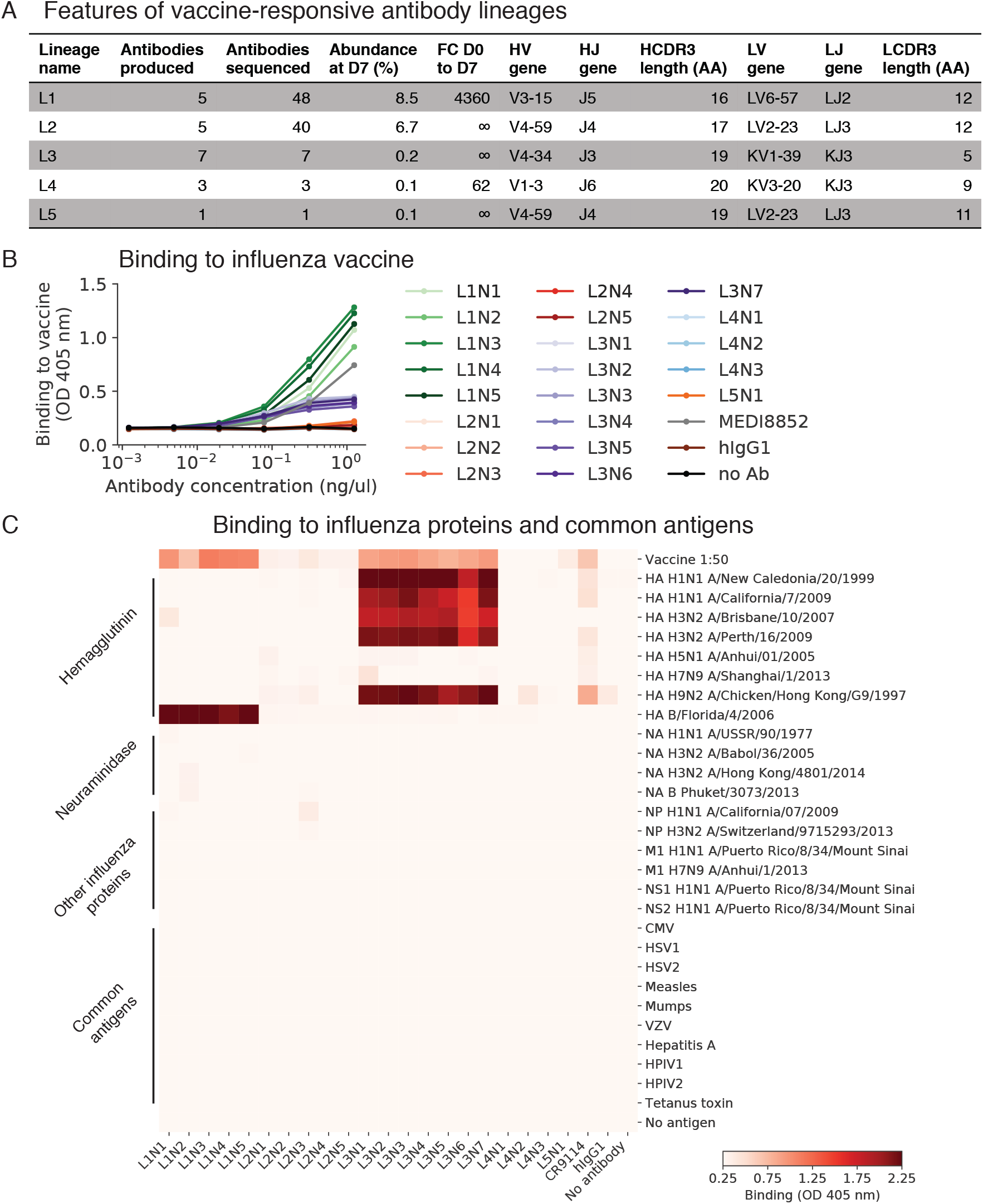
Molecular features and functional characterization of influenza vaccine-responsive antibodies. (**A**) Molecular features and population dynamics of vaccine-responsive antibody clones. FC, fold-change; D0, day 0 after vaccination; D7, day 7 after vaccination; AA, amino acids. (**B** and **C**) Binding of vaccine-responsive antibodies to the vaccine given to the subject (trivalent inactivated influenza vaccine from the 2011–2012 flu season) (**B**) and to purified influenza proteins and common viral and bacterial antigens (**C**) was measured using enzyme-linked immunosorbent assay (ELISA). HA, hemagglutinin; NA, neuraminidase; NP, nucleoprotein, M1, matrix protein 1; NS1, non-structural protein 1; NS2, non-structural protein 2; CMV, cytomegalovirus; HSV1/2, herpes simplex virus 1/2; VZV, varicella zoster virus; HPIV1/2, human parainfluenza virus 1/2; OD, optical density; hIgG1, human IgG1; no Ab, no antibody.

**Figure S4.**
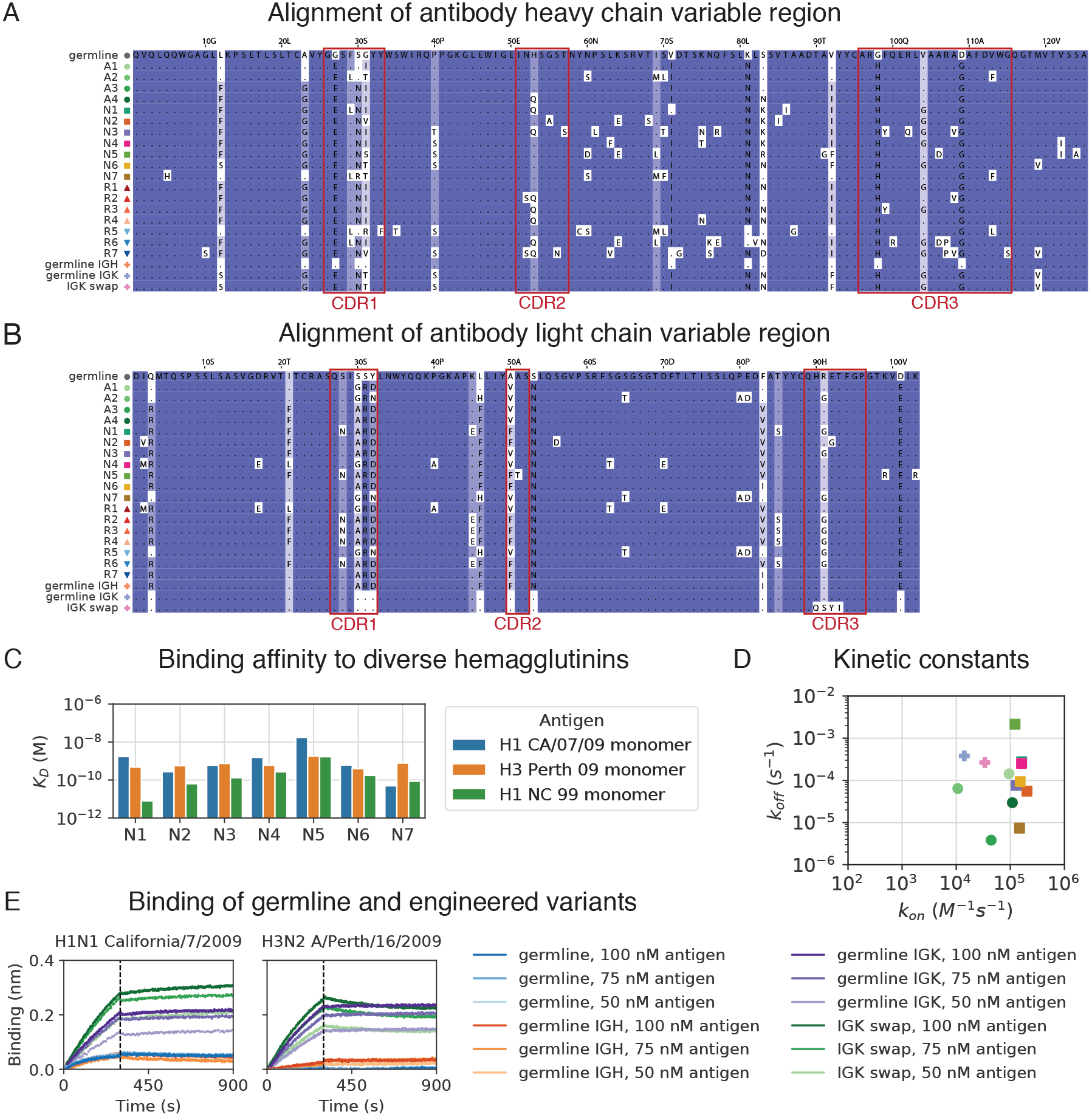
Molecular features and functional characterization of a broadly binding anti-influenza antibody clone. (**A** and **B**) Alignments of heavy (**A**) and light (**B**) chain variable region protein sequences for antibodies and engineered variants from clone L3. CDRs are indicated by red boxes. Background color indicates conservation of the position. Residues that are the same as germline are indicated by “.”. (**C** and **D**) Equilibrium constants (K_D_) (**C**) and kinetic constants (k_on_ and k_off_) (**D**) of binding between antibody variants from L3 and hemagglutinin variants were measured using biolayer interferometry. Symbols denoting variants are shown in Figure S4A. (**E**) Kinetics of binding and unbinding of germline and engineered antibody variants to H1 (A/California/7/2009) (left) and H3 (A/Perth/16/2009) (right) hemagglutinin antigens. Colors indicate antibody variants and antigen concentration. Dashed line indicates transition from association to dissociation step. Note that determination of equilibrium binding constants (shown in Figure 4C) was performed at lower antigen concentrations (not shown here).

**Figure S5.**
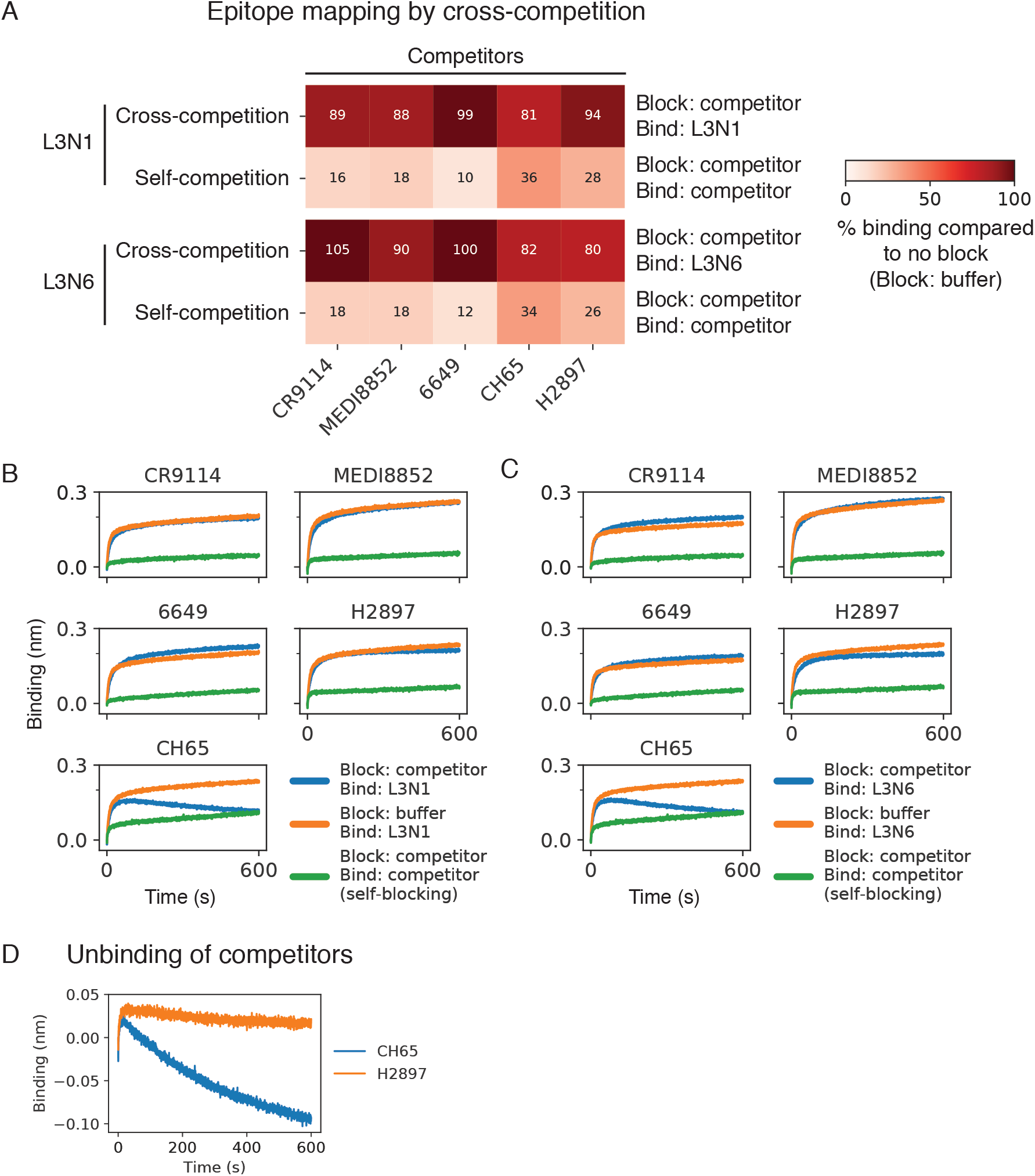
Determination of antibody epitopes by cross-competition. (**A**–**C**) Binding of antibodies L3N1 and L3N6 to trimeric hemagglutinin (H1 A/New Caledonia/20/1999) following blocking with potentially competing antibodies was determined using biolayer interferometry. To summarize the data (**A**), binding was determined after 50 s and compared with binding observed without blocking (using buffer instead of a potentially competing antibody during the blocking step). Numerical values are shown in each condition. Kinetics of binding are shown for L3N1 (**B**) and L3N6 (**C**). (**D**) Kinetics of unbinding for competitors CH65 and H2897 are shown. Fast unbinding of CH65 explains the observed decrease in binding in (**B** and **C**).

**Table S1. Genes that are differentially expressed between naïve and memory B cells.** Differential expression between single-cell transcriptional profiles of naïve (n = 18,953) and memory (n = 16,653) B cells was determined using the negative binomial exact test with the Benjamini-Hochberg correction for multiple testing. Genes with P < 0.05 are shown. Genes upregulated in memory B cells are shown first, then genes upregulated in naïve B cells.

**Table S2. Genes that are differentially expressed between memory and activated memory B cells.** Differential expression between single-cell transcriptional profiles of memory (n = 16,653) and activated memory (n = 421) B cells was determined using the negative binomial exact test with the Benjamini-Hochberg correction for multiple testing. Genes with P < 0.05 are shown. Genes upregulated in activated memory B cells are shown first, then genes upregulated in memory B cells.

